# Long-read, whole-genome shotgun sequence data for five model organisms

**DOI:** 10.1101/008037

**Authors:** Kristi E. Kim, Paul Peluso, Primo Babayan, P. Jane Yeadon, Charles Yu, William W. Fisher, Chen-Shan Chin, Nicole Rapicavoli, David R. Rank, Joachim Li, David E. A. Catcheside, Susan E. Celniker, Adam M. Phillippy, Casey M. Bergman, Jane M. Landolin

## Abstract

Single molecule, real-time (SMRT) sequencing from Pacific Biosciences is increasingly used in many areas of biological research including de novo genome assembly, structural-variant identification, haplotype phasing, mRNA isoform discovery, and base-modification analyses. High-quality, public datasets of SMRT sequences can spur development of analytic tools that can accommodate unique characteristics of SMRT data (long read lengths, lack of GC or amplification bias, and a random error profile leading to high consensus accuracy). In this paper, we describe eight high-coverage SMRT sequence datasets from five organisms (Escherichia coli, Saccharomyces cerevisiae, Neurospora crassa, Arabidopsis thaliana, and Drosophila melanogaster) that have been publicly released to the general scientific community (NCBI Sequence Read Archive ID SRP040522). Data were generated using two sequencing chemistries (P4C2 and P5C3) on the PacBio RS II instrument. The datasets reported here can be used without restriction by the research community to generate whole-genome assemblies, test new algorithms, investigate genome structure and evolution, and identify base modifications in some of the most widely-studied model systems in biological research.

## Background and Summary

Single-molecule, real-time (SMRT^®^) DNA sequencing occurs by optically detecting a fluorescent signal when a nucleotide is being incorporated by a DNA polymerase^1^. This relatively new technology enables detection of DNA sequences that have unique characteristics, such as long read lengths, lack of CG bias, random error profiles, and can yield highly accurate consensus sequences. Kinetic information such as pulse width and interpulse duration are also recorded and can be used to detect base modifications^2^,^3^.

Since its introduction, investigators have published on a range of applications using SMRT sequencing. For example, the developers of GATK (Genome Analysis Toolkit) demonstrated that single nucleotide polymorphisms (SNPs) could be detected using SMRT sequences^4^,^5^ due to their lack of context-specific bias and systematic error^5^,^6^. Likewise, the developers of PBcR (PacBio error correction)^7^,^8^ showed that complete bacterial genome assemblies using SMRT sequence data were achievable and had greater than Q60 consensus base quality^8^. PBcR was later incorporated as the “pre-assembly” step in the HGAP (hierarchical genome assembly process) system^9^, followed by consensus polishing using the Quiver algorithm^9^ to produce a complete assembly pipeline in SMRT Analysis, a free and open-source software suite released by Pacific Biosciences. In addition, other third-party tools now support long reads for various applications such as mapping^10^,^11^, scaffolding^12^, structural-variation discovery^13^, and genome assembly^7^,^14^. Other applications such as 16S rRNA sequencing^15^, characterization of entire transcriptomes in chickens^16^ and humans^17^, genome-editing studies^18^, base-modification studies^19^,^20^,^21^,^22^, and validation of CRISPR targets^23^ have also been published. Several datasets from this publication have already been used to develop the MinHash Alignment Process (MHAP), a new method for fast and efficient overlap of long reads for assembling large genomes^24^.

To encourage interest in further applications and tool development for SMRT sequence data, we report here the release of eight whole-genome shotgun-sequence datasets from five model organisms (*E. coli, S. cerevisiae, N. crassa, A. thaliana, and D. melanogaster*). These organisms have among the most complete and well-annotated reference genome sequences, due to continual refinement by dedicated teams of scientists. Despite continued improvement of these genome sequences with new technologies, few are completely finished with fully contiguous assemblies of all chromosomes. The gaps remaining arise from complex structures such as transposable elements, repeats, segmental duplications, or other dynamic regions of the genome that cannot be easily assembled. Structural differences in these regions can account for variability in millions of nucleotides within every genome, and mounting evidence suggest that such mutations are important for human diversity and disease susceptibility in many complex traits^25^ including autism and schizophrenia^26^. SMRT sequencing data can therefore play an important role in the completion of these and other reference genomes, providing a platform for new insights into genome biology.

## Methods

We generated eight whole-genome shotgun-sequence datasets from five model organisms using the P4C2 or P5C3 polymerase and chemistry combinations, totaling nearly 1000 gigabytes (GB) of raw data (See Data Records section). Genomic DNA was either purchased from commercial sources or generously provided by collaborators.

Genomic DNA sample summaries are provided in Table 1. DNA from the reference K12 strain of E. coli was purchased from Lofstrand Labs Limited (K12 MG1655 *E. coli*, cat# L3-4001SP2). DNA from the reference OR74A strain of *N. crassa* was purchased from the Fungal Genetics Stock Center (FGSC). A standard Ler-0 strain of *A. thaliana* plants was grown from seeds purchased from Lehle seeds (WT-04-19-02) and DNA was extracted at Pacific Biosciences. The protocol is available on Sample Net^27^ and summarized in the organism-specific methods section of this paper. DNA from the 9464 strain of *S. cerevisiae* was provided by J. Li at University of California San Francisco. The 9464 strain is a daughter of the reference WG303 strain. DNA from the T1 strain of *N. crassa* was obtained from D. Catcheside at Flinders University who has an interest in polymorphic genes regulating recombination. The T1 strain is an A mating type strain which, like OR74A, was derived from a cross between the Em a 5297 and Em A 5256 strains. DNA from the ISO1 strain^28^ of *D. melanogaster* was obtained from S. Celniker at Lawrence Berkeley National Laboratory. This is the reference strain of *D. melanogaster* that was originally chosen to be the first large genome to be sequenced and assembled using a whole-genome shotgun approach^29^. It continues to serve as the reference strain in subsequent releases and numerous annotations of the *D. melanogaster* genome.

**Table 1.** Summary of DNA Samples. The NCBI sample ID associated with each dataset is provided. DNA was extracted in a species-specific manner, yielding genomic DNA of various sizes. All DNA was size selected using the Blue Pippin system (Sage Sciences), and select samples were sheared with g-TUBEs (Covaris).

| Dataset Name | Sample ID | DNA extraction | gDNA size (kb) | Shearing | Size selection |
| --- | --- | --- | --- | --- | --- |
| *E. coli* MG1655 P4C2 | SAMN02951645 | ammonium acetate or SDS, proteinase K, phenol-chloroform | 17 | none | Blue Pippin (7kb) |
| *E. coli* MG1655 P5C3 | SAMN02743420 | ammonium acetate or SDS, proteinase K, phenol-chloroform | 17 | none | Blue Pippin (7kb) |
| *S. cerevisiae* 9464 P4C2 | SAMN02731377 | contact J. Li at UCSF | >40 | g-TUBE | Blue Pippin (17kb) |
| *N. crassa* OR74A P4C2 | SAMN02724975 | BashingBeads, Zymo Research kit | 6 | none | Blue Pippin (4kb) |
| *N. crassa* T1 P4C2 | SAMN02724976 | SDS, proteinase K, phenol-chloroform, RNAase, isopropanol | 15 | none | Blue Pippin (7kb) |
| *A. thaliana* Ler-0 P4C2 | SAMN02731378 | CTAB, chloroform:isoamyl, isopropanol precip. | >40 | g-TUBE | Blue Pippin (7kb) |
| *A. thaliana* Ler-0 P5C3 | SAMN02724977 | CTAB, chloroform:isoamyl, isopropanol precip. | >40 | g-TUBE | Blue Pippin (15kb) |
| *D. melanogaster* ISO1 P5C3 | SAMN02614627 | SDS, phenol-chloroform, CsCl banding, ethanol precip. | >40 | g-TUBE | Blue Pippin (17kb) |

DNA extraction methods were species-specific and optimized for each organism (see organism-specific methods below). In general, the steps are: (1) remove debris and particulate material, (2) lyse cells, (3) remove membrane lipids, proteins and RNA, (4) DNA purification.

SMRTbell^™^ libraries for sequencing^4^ were prepared using either 10 kb^30^,^31^ or 20 kb^32^ preparation protocols to optimize for the most high-quality and longest reads. The main steps for library preparation are: (1) shearing, (2) DNA damage repair, (3) blunt end-ligation with hairpin adapters supplied in the DNA Template Prep Kit 2.0 (Pacific Biosciences), (4) size selection, and (5) binding to polymerase using the DNA Sequencing Kit 3.0 (Pacific Biosciences).

### E. coli collection, DNA Extraction, and SMRTbell Library Preparation

Both P4C2 and P5C3 samples were prepared in the same way. *E. coli* K12 genomic DNA was ordered and purified by Lofstrand Labs Limited (K12MG1655 *E. coli*, cat# L3-4001SP2). Field Inversion Gel Electrophoresis (FIGE) was run to ensure presence of high-molecular-weight gDNA. Ten micrograms of gDNA was sheared using g-TUBE devices (Covaris, Inc) spun at 5500 rpm or 2029g on the MiniSpin Plus (Eppendorf) for 1 minute. Three microliters of elution buffer (EB) was added to rinse the upper chamber, spun at 6000 rpm, or 2415g and spun again at 5500 rpm or 2029g on the MiniSpin Plus (Eppendorf) after inverting the g-TUBE device. SMRTbell libraries were created using the “Procedure & Checklist – 20 kb Template Preparation using BluePippin™ Size Selection" protocol^32^. Briefly, the library was run on a BluePippin system (Sage Science, Inc., Beverly, MA, USA) to select for SMRTbell templates greater than 10 kb. The resulting average insert size was 17 kb based on 2100 Bioanalyzer instrument (Agilent Technologies Genomics, Santa Clara, CA., USA). Sequencing primers were annealed to the hairpins of the SMRTbell templates followed by binding with the P5 sequencing polymerase and MagBeads (Pacific Biosciences, Menlo Park, CA, USA). One SMRT Cell was run on the PacBio^®^ RS II system with an on-plate concentration of 150 pM using P5C3 chemistry and a 180-minute data-collection mode.

### S. cerevisiae collection, DNA Extraction, and SMRTbell Library Preparation

The 9464 strain is a MAT a haploid strain derived from a w303 reference strain following three integration events: 1) a construct (pTef2-dTomato-kanCMX6) was inserted near the CEN4 gene; 2) a construct (pTEF2-eGFP, natMX) was inserted near the ESP gene; and 3) the pds1 gene was deleted and replaced with the URA3 gene from *Kluyveromyces lactis*. Cells were grown to an OD600 of ~2 and 350 OD units of cells corresponding to roughly 7×10^9^ cells were harvested by centrifugation. Cells were washed in 4 ml of TE then resuspended in 4 ml of Buffer Y1 (Qiagen genomic DNA prep) and spheroplasted by addition of 250 units of Zymolyase 100T Seikagaku 120493) for 40 min at 30° C. Spheroplasts were pelleted and re-suspended in 5mL of Qiagen Buffer G2 containing 300 micrograms of Qiagen RNAse to lyse the cells. 2mg of proteinase K was then added to the lysate, which was incubated at 50° C for 30 min. The lysate was then centrifuged at 5000 G for 10 min at 4° C, and the supernatant was purified on a Qiagen 100/G genomic prep tip as per Qiagen instructions. The eluted DNA was spooled by addition of 3.5 ml of isopropanol to the 5 ml of eluated. The DNA was washed in 70% EtOH, air dried, and re-suspended in 200 microliters of TE by slowly dissolving overnight at room temperature. SMRTbell libraries were created using the “Procedure and Checklist – 10 kb Template Preparation and Sequencing (with Low-Input DNA)” protocol^30^. Twelve SMRT Cells were run on the PacBio RS II system using P4C2 chemistry and a 180-minute data collection mode.

### N. crassa OR74A, collection, DNA Extraction, and SMRTbell Library Preparation

*N. crassa* OR74A was purchased from FGSC# 2489. Cells were inoculated to a density of 4×10^6^ conidia in 75 ml Vogel's minimal medium (Medium N)^33^ and incubated at room temperature with gentle shaking for approximately 48 hours. Visual inspection shows the culture prior to harvest and demonstrates that there was only vegetative tissue and no asexual sporulation or induction of aerial hyphae. Mycelia was blotted dried on sterile paper toweling and pulverized for approximately 30 seconds at half of the maximum setting in a Biospec Products Mini BeadBeater tissue disruptor using the disrupting beads provided with the Zymo Research ZR fungal/bacterial DNA midi prep kit. Tissue was removed with a sterile inoculating stick and 100 mg of dried mycelia per sample was processed according to the manufactures instructions. DNA was eluted into 100 ul sterile water and DNA from two samples was pooled. Yield was quantified using a nano drop system and also validated by agarose gel electrophoresis. The concentration was 32.57 ng/ul, A260 was 0.651, A280 was 0.373, 260/280 was 1.75 and 260/230 was 0.91. The genomic DNA was approximately 6 kb and was not sheared. SMRTbell libraries were created using the “Procedure and Checklist – 10 kb Template Preparation and Sequencing (with Low-Input DNA)” protocol^30^. Two SMRT Cells were run on the PacBio RS II system using P4C2 chemistry and a 180-minute data collection mode.

### N. crassa T1 collection, DNA Extraction, and SMRTbell Library Preparation

The T1 strain of *N. crassa*, is an A mating type strain derived by DG Catcheside from a cross between the Em a 5297 and Em A 5256 strains he obtained from Stirling Emerson in 1955. The fungus was grown in shake culture for 72 hr at 25°C in 500 ml Vogel's N ^34^ minimal medium containing 2% sucrose. Mycelium was harvested by filtration, ground in liquid nitrogen, resuspended in 10 ml of a buffer containing 0.15 M NaCl, 0.1 M EDTA, 2% SDS at pH 9.5, and incubated overnight at 37°C with 1 mg protease K. Debris was precipitated by centrifugation and 10 ml distilled water was added to the supernatant, which was extracted once with an equal volume of water-saturated phenol and once with chloroform. Nucleic acids were precipitated from the aqueous phase with 0.6 volumes of isopropanol. Following centrifugation, the pellet was dried and dissolved in 1 ml TE buffer (TRIS 10 mM, 1 mM EDTA pH 8.0). RNA and protein were digested by overnight incubation at 37°C with RNAase (50 µg) followed by addition of protease K (50 µg) and further incubation for 2 hr. The digest was extracted once with water-saturated phenol and once with chloroform. DNA was collected by precipitation with 0.6 volumes of isopropanol and, following centrifugation, the pellet was dried, dissolved in 500 µl TE buffer and stored at 4°C. Field Inversion Gel Electrophoresis (FIGE) was run to ensure presence of high-molecular-weight gDNA. The genomic DNA was approximately 15 kb and was not sheared. SMRTbell libraries were created using the “Procedure and Checklist – 10 kb Template Preparation and Sequencing (with Low-Input DNA)” protocol^30^. Eighteen SMRT Cells were run on the PacBio RS II system using P4C2 chemistry and a 180-minute data-collection mode.

### A. thaliana collection, DNA Extraction, and SMRTbell Library Preparation

Plants were grown from seeds provided by Lehle seeds (WT-04-19-02). Shoots and leaves were harvested at three weeks and ground in liquid nitrogen using a mortar and pestle. The complete protocol is described in the “Preparing *Arabidopsis* Genomic DNA for Size-Selected ∼20 kb SMRTbell^™^ Libraries” protocol^35^. This protocol can be used to prepare purified *Arabidopsis* genomic DNA for size-selected SMRTbell templates with average insert sizes of 10 to 20 kb. We recommend starting with 20-40 grams of three-week-old *Arabidopsis* whole plants, which can generate >100 μg of purified genomic DNA. SMRTbell libraries were created using the “Procedure & Checklist – 20 kb Template Preparation using BluePippin^™^ Size Selection” protocol^32^. Eighty-five SMRT Cells were run on the PacBio RS II system using P4C2 chemistry and a 180-minute data-collection mode. Forty-six SMRT Cells were run on the PacBio RS II system using P5C3 chemistry and a 180-minute data-collection mode.

### D. melanogaster collection, DNA Extraction, and SMRTbell Library Preparation

A total of 1.2 g of adult male ISO1 flies corresponding to 1950 animals were collected, starved for 90-120 min and frozen. The flies ranged in age from 0-7 days based on four collections (1) 0-2 days old, 500 males, 0.33 g; (2) 0-4 days old, 500 males, 0.29 g; (3) 0-7 days old, 500 males, 0.29 g; (4) 0-2 days old, 450 males, 0.29 g. Flies were ground in liquid nitrogen to a fine powder and genomic DNA was purified by phenol-chloroform extraction and CsCl banding in the ultracentrifuge. Briefly, the pulverized fly extract was gently re-suspended in 15 ml of HB buffer (7 M Urea, 2% SDS, 50 mM Tris pH7.5, 10 mM EDTA and 0.35 M NaCl) and 15 ml of 1:1 phenol/chloroform. The mixture was shaken slowly for 30 minutes and then centrifuged at 23,600g for 10 min at 20°C in a Sorvall HB-4 rotor. The aqueous phase was re-extracted twice as above and then precipitated by adding two volumes of ethanol and centrifuging at 23,600g for 10 min at 20°C in a Sorvall HB-4 rotor. The pellet was re-suspended in 3 ml of TE (10 mM Tris 1 mM EDTA pH 8.0) by gentle inversion for 3 hrs. To the re-suspended DNA, 3 g CsCl and 0.3 ml of 10 mg/ml ethidium bromide (EtBr) were added and the mixture centrifuged at 199,000g for 16 hrs at 15°C in a Beckman VTi 65.2 rotor. The EtBr was removed by extraction with water-saturated butanol, performed 3 times in a Beckman JA-12 rotor at 13,000g for 5 min. at 4 degree C each time. The DNA was diluted three-fold with TE, 1/10 vol, 4 M NaCl was added and the DNA precipitated with two volumes of ethanol. Centrifugation was done in a JA-12 rotor at 16,000g for 30 min at 4 degree C for the precipitation step. After centrifugation, the pellet was washed in 70% ethanol. The DNA was resuspended in 100 μl TE at a concentration of 1.4 μg/μl and quantified using a Nanodrop instrument. This protocol routinely yields at least 10 ng DNA per mg of flies with an estimated DNA size >100 kb.Genomic DNA was sheared using a g-TUBE device (Covaris) at 4800 rpm or 1545 g on the MiniSpin Plus (Eppendorf), 150 ng/μl and purified using 0.45x volume ratio of AMPure PB beads. SMRTbell libraries were created using the Procedure & Checklist – 20 kb Template Preparation using BluePippin^™^ Size Selection^32^. Libraries were ligated with excess adapters and an overnight incubation was performed to increase the yield of ligated fragments larger than 20 kb. Smaller fragments and adapter dimers were then removed by >15 kb size selection using the BluePippin DNA size selection system by Sage Science. Forty-two SMRT Cells were run on the PacBio RS II system. The first run was composed of four SMRT Cells, loaded at 75 pM, 150 pM, 300 pM, and 400 pM in order to determine the optimal loading concentration of the sample. The remaining 38 SMRT Cells were loaded at 400 pM.

## Data Records

After DNA extraction, libraries were generated and sequenced at Pacific Biosciences of California, uploaded to Amazon Web Services' Simple Storage Service (S3), and then submitted to the Sequence Read Archive (SRA) at NCBI under Project ID SRP040522 (Data Citation 1). The corresponding accession numbers and file sizes are listed in Table 2. More detailed information including md5 checksums and links to download the original data from AWS S3 are provided in Supplementary Table S1.

**Table 2.** Summary of Datasets. Eight datasets from five organisms are described in this paper. Data can be accessed from SRA using the accession numbers provided.

| Organism | Strain | Origin | Polymerase & Chemistry Library kits | SRA Accession | Size (GB) |
| --- | --- | --- | --- | --- | --- |
| *E. coli* | MG1655 | Lofstrand Labs | P4C2 | SRX669475 | 6.0 |
| *E. coli* | MG1655 | Lofstrand Labs | P5C3 | SRX533603 | 3.8 |
| *S. cerevisiae* | 9464 | J. Li | P4C2 | SRX533604 | 38 |
| *N. crassa* | OR74A | FGSC | P4C2 | SRX533605 | 29 |
| *N. crassa* | T1 | D. Catcheside | P4C2 | SRX533606 | 143 |
| *A. thaliana* | Ler-0 | Lehle Seeds | P4C2 | SRX533608 | 263 |
| *A. thaliana* | Ler-0 | Lehle Seeds | P5C3 | SRX533607 | 252 |
| *D. melanogaster* | ISO1 | S. Celniker | P5C3 | SRX499318 | 187 |

Raw data was transferred from the instrument to a storage location and organized first by the run name, and then by the SMRT Cell directory. Each run contained one or more SMRT Cells. Each SMRT Cell produced a metadata.xml file that recorded the run conditions and barcodes of sequencing kits, three bax.h5 files that contained base call and quality information of actual sequenced data, and one bas.h5 file that acted as a pointer to consolidate the three bax.h5 files. The “h5” suffix denotes that these are Hierarchical Data format 5 (HDF5) files. The specific contents and structure of a PacBio bax.h5 file is described in more detail in online documentation^36^.

Recall the “SMRT bell” structure that underwent sequencing was created by the library preparation process ^4^. Sequenced SMRT Bells corresponded to raw reads that may pass around the same base multiple times. A raw read could therefore have a structure that is composed of left adapter→ DNA insert → right adapter → reverse complement of DNA insert → left adapter → DNA insert → and so on. This raw read is typically processed downstream to remove adapters and create subreads composed of the DNA sequence of interest to the investigator. See the Technical Validation section for details on filtering parameters and free software used to analyze and quality check the data.

All datasets were filtered and mapped using SMRT Analysis v2.0.1, v2.1.0, or v2.2. There are no changes to the filtering or mapping parameters in these versions, and detailed parameters used are discussed in the Technical Validation section. SMRT Analysis is a software suite that is free and can be downloaded from the Pacific Biosciences Developer's Community Network Website (DevNet)^37^. SMRT Analysis includes the SMRT Portal graphical user interface, as well as SMRT View genome browser. Extensive documentation and technical support can also be accessed via the DevNet website.

The post-filter statistics of each dataset are listed in Table 3. While read lengths reflect the true sequencing capacity of the instrument, only subreads are summarized in Table 3 because it is relevant and used for downstream analysis algorithms such as *de novo* assemblers. Multiple subreads can be contained within one raw read, and subreads exclude adapters and low quality sequence. N50 is a statistic used to describe the length distribution of a collection of reads, contigs, or scaffolds, and is defined as the length where 50% of all bases are contained in sequences longer than that length. The N50 filtered subread lengths ranged from 7.6 kb to 10.5 kb for datasets generated with P4C2 chemistry and ranged from 12.2 kb to 14.2 kb for datasets generated with P5C3 chemistry. With the exception of *N. crassa* OR74A, all datasets were sequenced to high-coverage (>68X) and sufficient for *de novo* genome assembly applications. The *N. crassa* OR74A dataset was sequenced to 25X coverage and should be sufficient for mapping, consensus SNP calling, and testing other applications.

**Table 3.** Summary statistics of filtered data. Results shown for each dataset are based on output of SMRT Portal analysis using the default filtering parameters (see text for details). Fold coverage is calculated relative to the estimated genome size.

| Dataset Name | Number of filtered subreads | N50 filtered subread length (nt) | Maximum filtered subread length (nt) | Total filtered subread (nt) | Estimated genome size (Mb) | Fold coverage |
| --- | --- | --- | --- | --- | --- | --- |
| *E. coli* MG1655 P4C2 | 61,019 | 7,586 | 22,609 | 331,516,965 | 5 | 66X |
| *E. coli* MG1655 P5C3 | 43,063 | 12,041 | 28,647 | 373,874,428 | 5 | 75X |
| *S. cerevisiae* 9464 P4C2 | 269,145 | 8,821 | 30,164 | 1,597,871,118 | 12 | 133X |
| *N. crassa* OR74A P4C2 | 175,926 | 7,617 | 30,845 | 981,884,113 | 40 | 25X |
| *N. crassa* T1 P4C2 | 210,480 | 10,462 | 36,227 | 11,497,185,440 | 40 | 287X |
| *A. thaliana* Ler-0 P4C2 | 1,338,320 | 8,769 | 41,753 | 8,129,670,483 | 120 | 68X |
| *A. thaliana* Ler-0 P5C3 | 2,067,212 | 12,188 | 47,445 | 17,714,447,516 | 120 | 148X |
| *D. melanogaster* ISO1 P5C3 | 1,561,929 | 14,214 | 44,766 | 15,194,174,294 | 160 | 95X |

## Technical Validation

### DNA and Sample Preparation

To assess the quality of genomic DNA received, we used Qbit (Life Technologies) and Nanodrop (Thermo Scientific) to measure the concentration of genomic DNA. Ideal samples had similar concentration estimates on both platforms, with A_230/260/230_ ratios close to 1:1.8:1, corresponding to what is expected of pure DNA. All samples presented here passed this screening criterion.

Next we assessed the size of the genomic DNA received. For genomic DNA where the size range was less than 17kb, we used the Bioanalyzer 21000 (Agilent) to determine the actual size distribution. For genomic DNA where the size range was greater than 17kb, we opted for pulse field gel electrophoresis to better estimate the larger size distributions. The sizes of the genomic DNA for each sample are listed in Table 1.

To ensure that the library insert sizes were in the optimal size range, we sheared genomic DNA using gTubes if the apparent size was greater than 40 kb. Alternatively, if the size was less than 40kb, then the DNA was not sheared and carried straight through to library preparation. Extremely small fragments (<100bp) and adapter dimers are eliminated by Ampure Beads. Adapter dimers (0-10bp) and small inserts (11-100bp) represented less than 0.01% of all the reads sequenced in all datasets. We additionally use the Blue Pippin (Sage Science) to ensure that the libraries had a physical size of 10kb or greater. The size cutoffs used for each sample are listed in Table 1.

### Analysis and Quality Filtering

To assess the quality of the libraries sequenced, we examined the percent of bases filtered by a standard QC procedure. Filtering conditions for high-quality SMRT sequence data are read score > 0.8, read length > 500 nt, subread length > 500 nt. In addition, the ends of reads are trimmed if they are outside of high-quality (HQ) regions, and adapter sequences between subreads are removed. All samples retained 71-97% of the bases after filtering. High-quality regions are defined by the base caller in primary analysis (on the PacBio RS II instrument) and indicate the contiguous region of the trace that contains high quality sequence data^38^. All datasets were filtered with the parameters above using SMRT Analysis v2.0.1, v2.1.0, or v2.2.0. There are no changes to the filtering protocol in these versions.

To ensure that the sequences matched the model organism of interest, we examined the percent of post-filter bases that were mapped to the closest reference genome available. All datasets were mapped using blasr^11^ from SMRT Analysis v2.2.0. Plots showing the distributions of mapped subread concordances are provided in Figure 1, and are a rough measure of how well the reads agree with the reference genome. Note that these numbers are an underestimate of the true accuracies of the reads because the DNA was not always from the same strain as the reference genome, and new stock may have evolved such that certain bases or structural repeats are different from the reference strain that was first sequenced decades ago. The depth of coverage for each chromosome is also plotted in figure 1.

**Figure 1.**
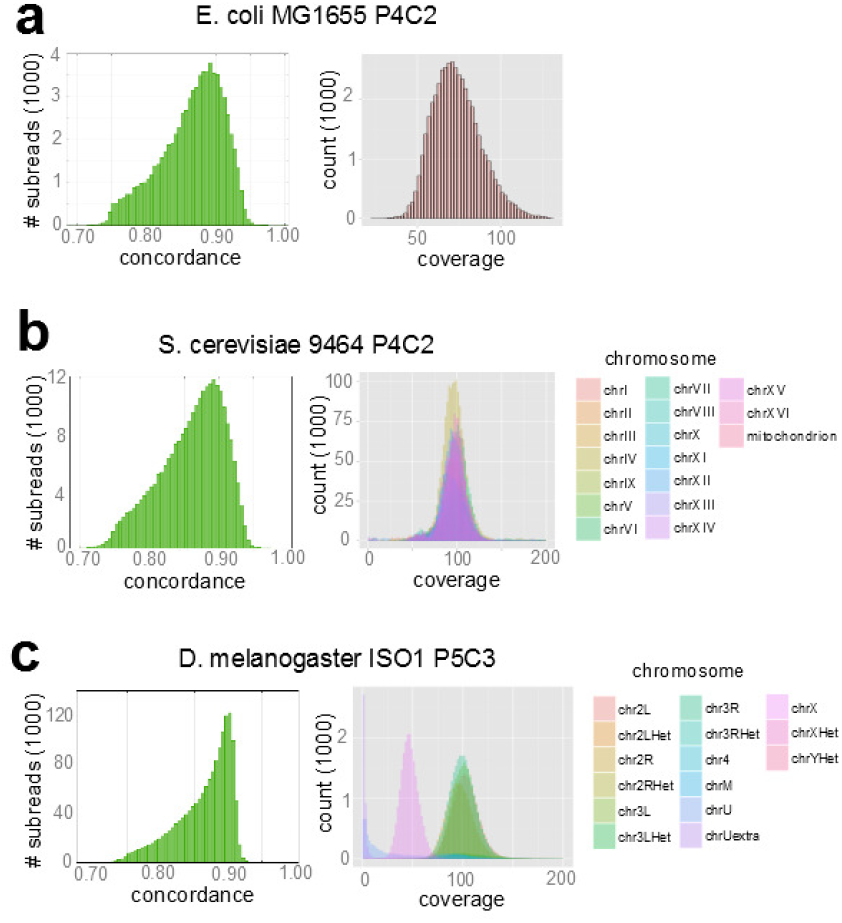
Mapped Subread Concordance and Coverage. The distribution of mapped subread concordances and mapped subread coverages are plotted for *E. coli* MG1655 P4C2 **(a)**, *S. cerevisiae* 9464 P4C2 **(b)**, and *D. melanogaster* ISO1 P5C3 **(c)**. The coverage distribution is similar among all chromosomes in *S. cerevisiae*, whereas the coverage distribution is half in chrX (50X) compared to the autosomes (100X) in *D. melanogaster*. ChrU and chrUextra are assembled contigs that could not be placed to physical chromosomes, and have very low coverages in general.

We used reference genomes that were from the closest strain that was available in the public domain. For *E. coli*, we used the sequence from NC_000913 at NCBI GenBank, which was first sequenced by Blattner *et al*.^39^ (Data Citation 2). For *S. cerevisae*, we downloaded release R64-1-1 of the S288c reference from the *Saccharomyces* Genome Database Project (http://downloads.yeastgenome.org/sequence/S288C_reference/genome_releases/)(Data Citation 3). The changes and updates to the reference genome are reviewed by Engel *et al*.^40^ For *N. crassa*, we downloaded the genome sequence from the *N. crassa* database hosted by the Broad institute, which uses the same sequences contained in the AABX00000000.3 project at NCBI GenBank (Data Citation 4). This data was from the OR74A strain and first sequenced by Galagan *et al*.^41^. For *A. thaliana*, we used the reference sequences from The *Arabidopsis* Information Resource^42^,(Data Citation 5) (TAIR) version 10 (ftp://ftp.arabidopsis.org/home/tair/Sequences/whole_chromosomes/), which was originally sequenced and analyzed by The *Arabidopsis* Genome Initiative^43^. For *D. melanogaster*, we used sequences from RELEASE 5 of the *Drosophila* reference genome downloaded from the Berkeley *Drosophila* Genome Project website (http://www.fruitfly.org/sequence/release5genomic.shtml). This data was from the ISO1 strain and has been updated since the referenced RELEASE 3 version by Celniker *et al*.^29^ (Data Citation 6).

All samples had a mapping rate of 81-95%, with the exception of the *Neurospora* T1 sample that had a mapping rate of 62%. This sample may have some damaged DNA as it had been stored in a freezer for over 20 years. Nonetheless, preliminary unpublished results show that the sequence from the *Neurospora* T1 sample can be successfully assembled into a genome that is more contiguous than the existing reference genome for *Neurospora*^44^ (http://figshare.com/articles/ENCODE_like_study_using_PacBio_sequencing/928630).

## Usage Notes

The eight datasets are available for download in two locations: (1) Amazon S3 repositories, which contain primary analysis data in the original formats provided by the PacBio RS II instrument (*.metadata.xml, *.bas.h5, & *.bax.h5 files), and (2) Sequence Read archive entries, which contain unfiltered subread base calls in sra format and can be converted to unfiltered subread fasta and fastq files using the SRA toolkit. Links to Amazon S3 repositories as well as SRA accession numbers are provided in Supplementary Table S1. While fastq files can be used to assess basic read characteristics and analyzed by most third-party tools, original formats are still needed for more sophisticated analyses using SMRT Portal or PacBio-specific algorithms. Download the primary data from Amazon S3 if applications such as Quiver consensus base calling or base modification analyses are desired. These analyses require additional information encoded within .bax.h5 files such as quality values, pulse width, and inter-pulse duration. While bax.h5 files can also be converted to fasta or fastq formats, download data from the SRA if sra, fasta, or fastq formatted files are desired.

The sequence IDs provided in the original formats (bax.h5 files) are different from those provided by the SRA in sra or fastq formats. The sequence IDs in the original formats contain information about the sequencing run itself. The date, time and instrument id is tracked by a “m” prefix; the SMRT Cell barcode, 8 pack number, and other information is tracked by a “c” prefix; and the “s” and “p” prefixes are now deprecated. For example, a subread with the ID `“m130227_130322_42141_c100505662540000001823074808081362_s1_p0/234/3_13049”` indicates that the sequencing run was started in Feburary 27^th^ 2013 (m130227) at 1:03:22PM (130322) on instrument number 42141, using SMRT Cell ID c100505662540000001823074808081362. This subread also originates from zero mode wave guide (ZMW) number 234 on the SMRT cell, and corresponds to bases 3 to 13049 of the raw read in the ZMW. The same read will be re-named by SRA to an arbitrary index prefixed by the SRR accession number, and the header line in the fasta file will, for example, appear as “>SRR1284620.6 length=13046” in the files downloaded from the SRA.

The datasets described in this paper were first released on DevNet^37^, the PacBio Software Developer Community Network website, with brief descriptions on the PacBio blog. DevNet typically hosts open-source software, while SampleNet^27^, the PacBio Sample Preparation Community Network website, typically hosts protocols for DNA extraction and library preparation. These websites provide valuable data and documentation about the technology, but are not considered a part of the traditional academic record. This Data Descriptor in Scientific Data provides an opportunity to describe the methodology and characteristics of the eight datasets in more detail and creates a citable entity for the scientific community.

DNA sequencing instruments and chemistries change rapidly, and PacBio SMRT sequencing is no exception. The datasets presented here are from P4C2 and P5C3 polymerase-chemistry combinations, spanning release dates from late-2013 to early-2014. These datasets represent some of the longest read lengths to date for these chemistries, and can be used to benchmark and develop new algorithms and the state of the art as the technology evolves.

## Acknowledgements

The contributions of AMP were funded under Agreement No. HSHQDC-07-C-00020 awarded by the Department of Homeland Security Science and Technology Directorate (DHS/S&T) for the management and operation of the National Biodefense Analysis and Countermeasures Center (NBACC), a Federally Funded Research and Development Center. The views and conclusions contained in this document are those of the authors and should not be interpreted as necessarily representing the official policies, either expressed or implied, of the U.S. Department of Homeland Security. In no event shall the DHS, NBACC, or Battelle National Biodefense Institute (BNBI) have any responsibility or liability for any use, misuse, inability to use, or reliance upon the information contained herein. The Department of Homeland Security does not endorse any products or commercial services mentioned in this publication. CMB was supported by Human Frontier Science Program Young Investigator grant RGY0093/2012.

We thank J. Korlach and E. Hauw for assistance in manuscript preparation, R. Stainer for *Neurospora* T1 sample preparation, and J. Trow at NCBI for assistance with data submission.

## Author contributions

KEK prepared libraries, sequenced, and analyzed data for the *N. crassa* OR74A, *N. crassa* T1, and *D. melanogaster* datasets.

PP grew plants from seed, prepared libraries, and sequenced DNA for the *A. thaliana* P4C2 and *A. thaliana* P5C3 datasets. He also sequenced DNA for the *S. cerevisae* 9464 dataset.

PB prepared libraries and sequenced the *E. coli* datasets.

PJY provided DNA for the *N. crassa* T1 dataset.

CY extracted DNA for the *D. melanogaster* dataset.

WF collected male flies for the *D. melanogaster* dataset.

CSC analyzed data.

NAR extracted DNA, prepared libraries, and coordinated the *S. cerevisae* 9464 dataset.

DRR grew plants from seed, extracted DNA and coordinated the *A. thaliana* P4C2 and P5C3 datas000ets.

JL extracted DNA and prepared libraries for the *S. cerevisae* 9464 dataset.

DC provided DNA for the *N. crassa* T1 dataset.

SEC extracted DNA and coordinated the *D. melanogaster* dataset.

AMP analyzed data, coordinated the project, and prepared the manuscript.

CMB analyzed data, coordinated the project, and prepared the manuscript.

JML deposited data to the SRA, analyzed data, coordinated the project, and prepared the manuscript.

## Data Citations

1. *NCBI Sequence Read Archive,* SRP040522 (2014).
2. *Genbank*, NC_000913 (2006).
3. *NCBI Assembly,* GCF_000146045.2 (2011).
4. *NCBI Assembly,* AABX00000000.3 (2013).
5. *NCBI Assembly,* GCF_000001735.3 (2011).
6. *NCBI Assembly,* GCF_000001215.2 (2007).

